## Supplementary figures S1-8, tables S1-3 for "Combinatorial CRISPR screen reveals *FYN* and *KDM4* as targets for synergistic drug combination for treating triple negative breast cancer"

Timothy K. Lu, 500 Technology Square, Cambridge MA 02139, USA.

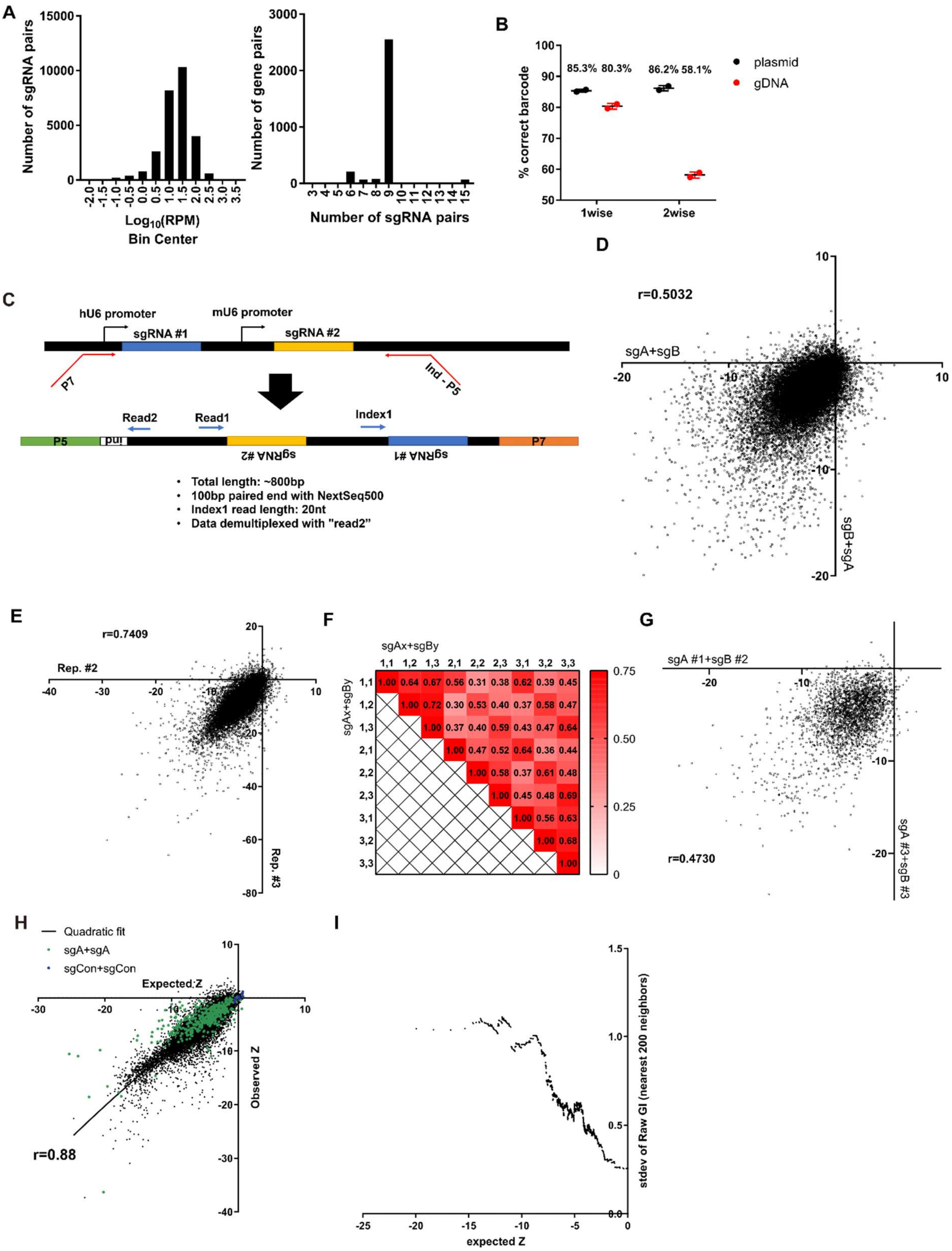


**Figure S1. Quality controls for combinatorial sgRNA libraries and screening results.** (A) Summary of representations of each sgRNA pair in the combinatorial library. (Left) Histogram of the frequency of each sgRNA combinations. (Right) Histogram of the number of sgRNA pairs that target a gene pair. (B) Accuracy of sgRNA-barcode pairing in each library (n=2). (C) Sequencing scheme for identifying both sgRNAs in the library by Next Generation Sequencing. (D) Scatter plot of growth phenotype score Z of two different permutations of the same sgRNA pairs. (E) Scatter plot of growth phenotype score Z of two independent biological replicates. (F) Heatmap of the Pearson’s r values between growth phenotype score of each sgRNA pairs that target the same gene pair. (G) Scatter plot of growth phenotype score Z of different sgRNA pairs that target the same gene pair. (H) Expected and Observed growth phenotype score Z of all sgRNA pairs. (I) Standard deviation of the GI score of 200 neighboring gene pairs by expected growth phenotype score. All data are plotted as mean±s.d. All replicates are biological replicates.

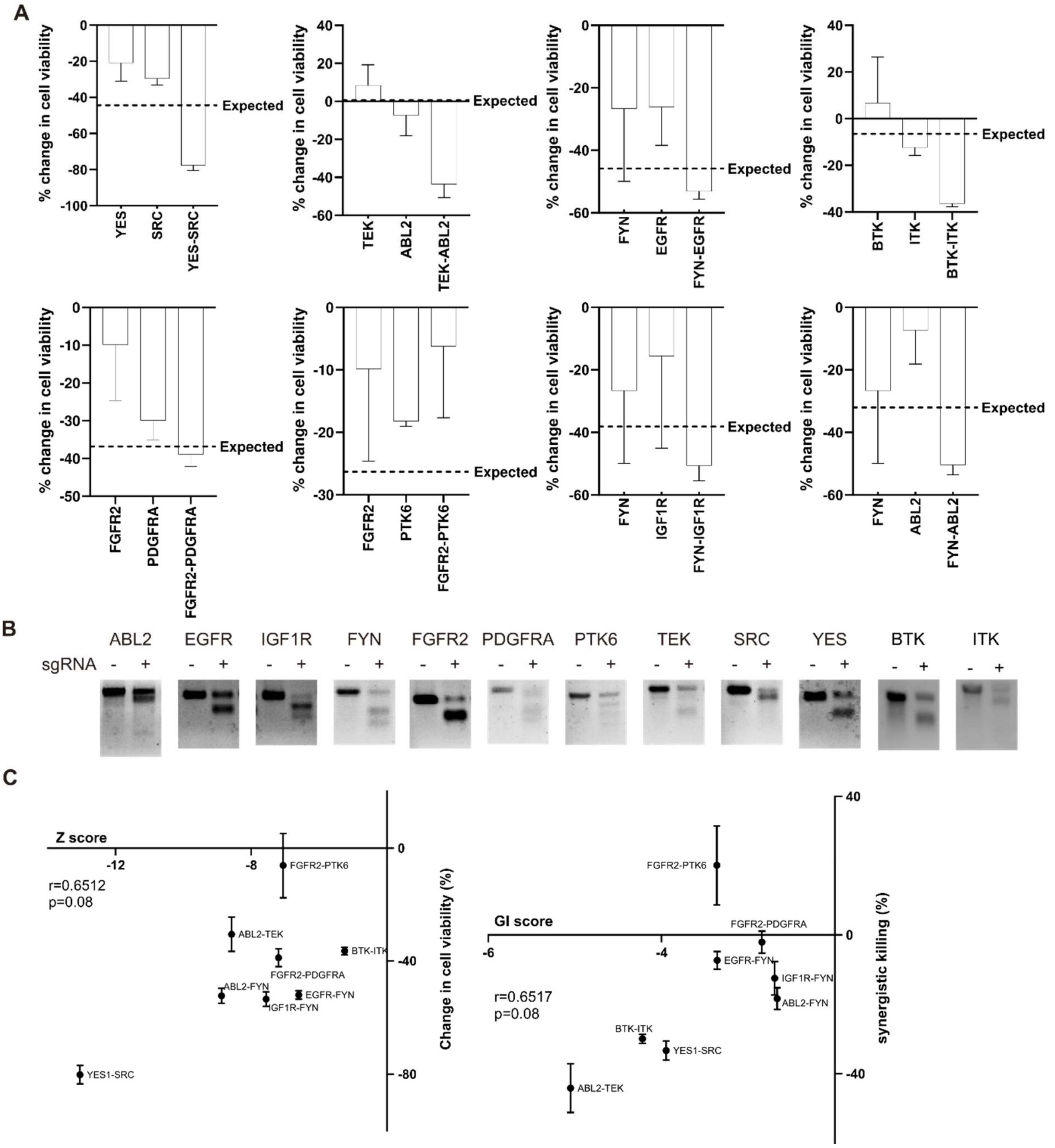


**Figure S2. Validation of screening hits.** (A) The changes in cell viability upon each single or double tyrosine kinase knockout in MDA-MB-231 cells analyzed in figure 2A-B. Expected double knockout phenotype was calculated using Bliss independence model. (B) T7 endonuclease assay of MDA-MB-231 cells infected with lentivirus expressing sgRNA targeting the indicated genes. (C) Scatter plot of (Left) growth phenotype score Z and the change in cell viability and (Right) GI score and synergistic killing by the indicated sgRNA combinations. Correlations were analyzed by Pearson's r. All data are plotted as mean±s.d. All replicates are biological replicates.

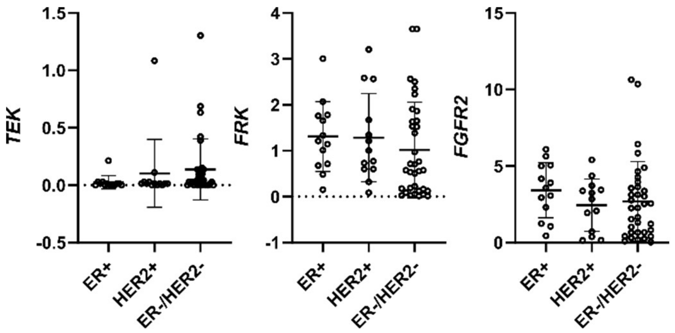


**Figure S3. Expression of TEK, FRK and FGFR2 in breast cancer cell lines of different subtypes.**

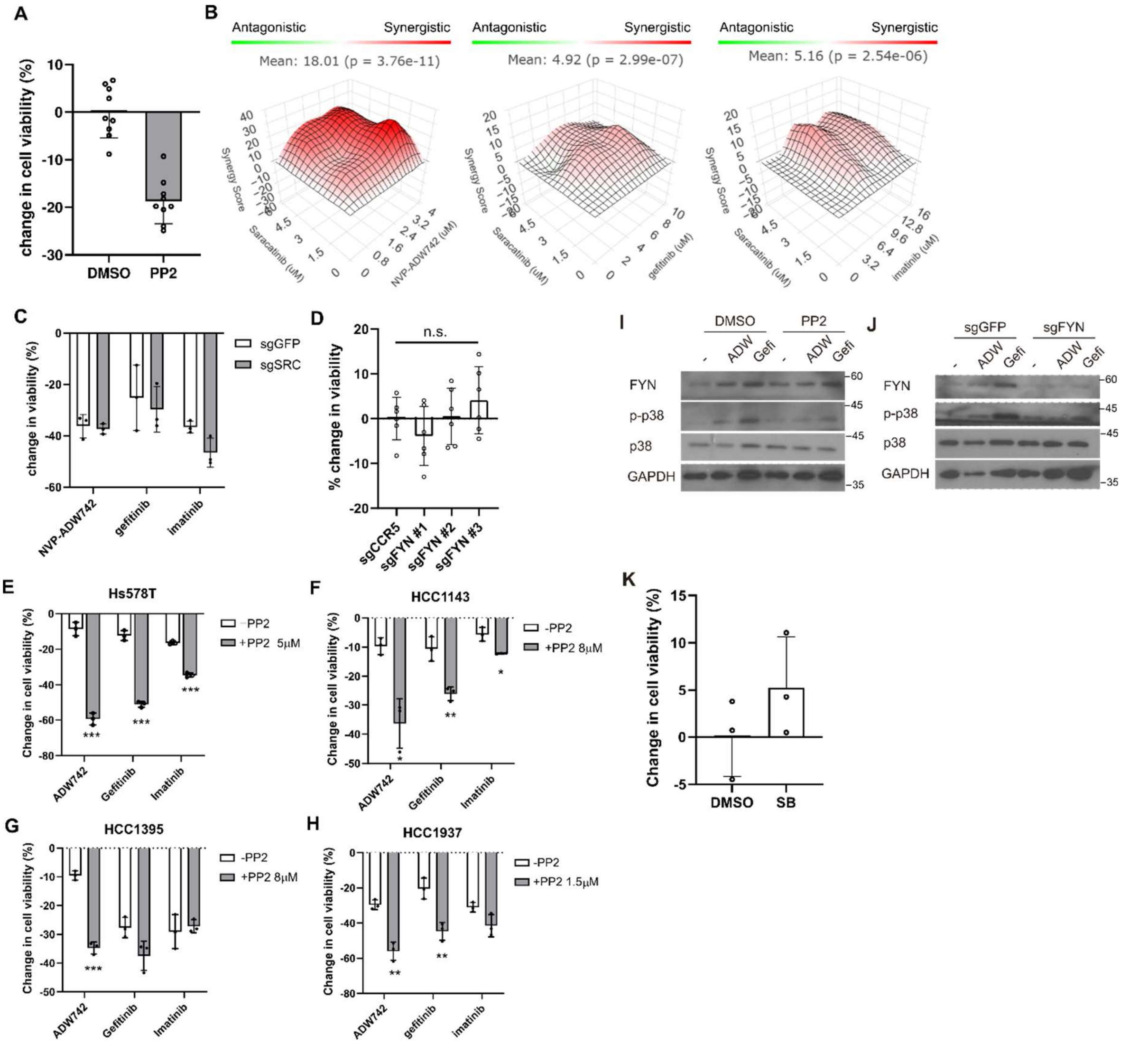


**Figure S4. FYN inhibition synergizes with TKIs in multiple TNBC cell lines.** (A) MTT assay with MDA-MB-231 cells treated with 10μM PP2 for 48 hours. (B) Synergistic killing calculated using SynergyFinder plus by Bliss independence model. (C) MTT assay with MDA-MB-231 Cas9 cells expressing indicated sgRNAs and treated with indicated TKIs (n=3). (D) MTT assay with MDA-MB-231 Cas9 cells expressing indicated sgRNAs. sgCCR5 is used as a control sgRNA that is expected to minimally affect cell viability. (E-H) MTT assay with various TNBC cell lines treated with indicated drug combinations (n=3). (I) western blot analysis of Hs578T cells treated with indicated drugs. (J) western blot analysis of Hs578T Cas9 cells expressing indicated sgRNA and treated with indicated drug. (K) MTT assay with MDA-MB-231 cells treated with 10μM SB203580 for 48 hours. All data are plotted as mean±s.d. Unpaired two-sided Student's t-test in D-H.*, $p<0.05$; **, $p<0.01$; ***, $p<0.001$; n.s., $p>0.05$. All replicates are biological replicates.

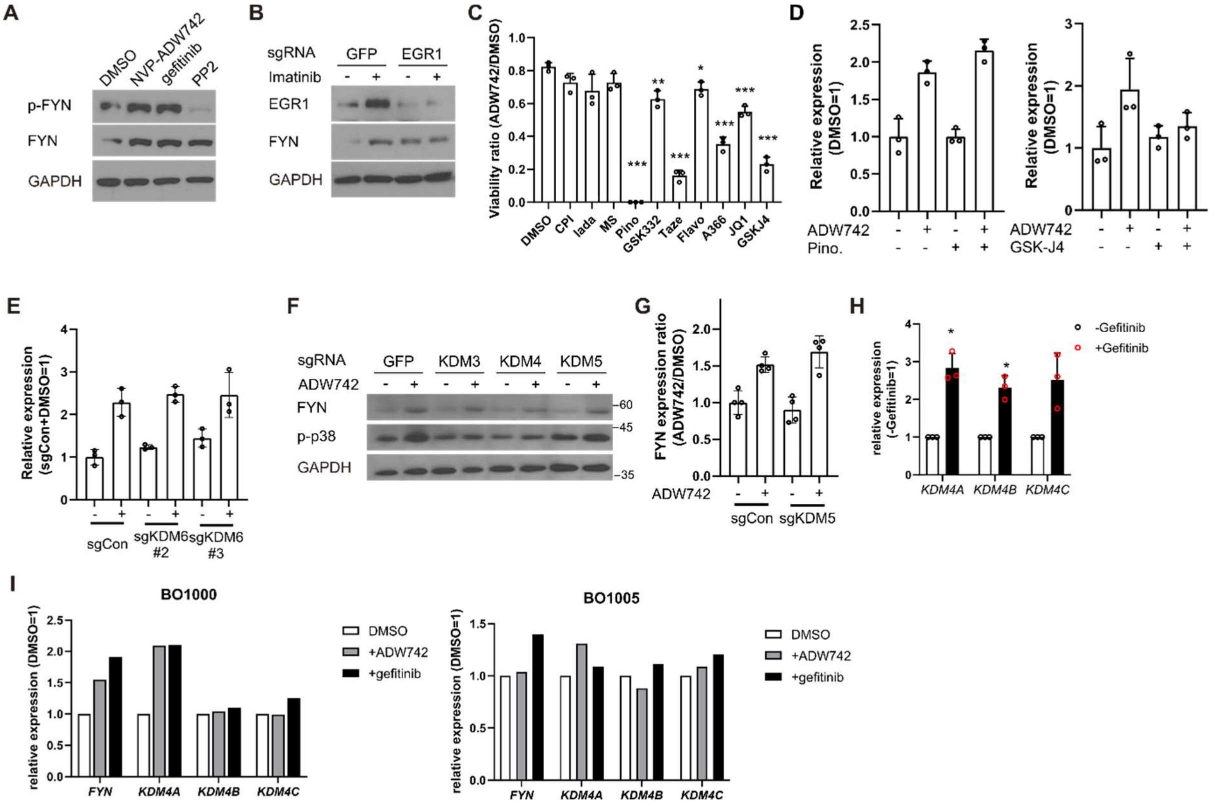


**Figure S5. KDM4 regulates FYN expression level, sensitizing MDA-MB-231 cells to TKIs.** (A) Western blot analysis of MDA-MB-231 cells treated with 4μM NVP-ADW742, 10 μM gefitinib, or 10 μM PP2 for 48 hours. (B) Western blot analysis of MDA-MB-231 Cas9 cells expressing indicated sgRNAs and treated with imatinib. (C) MTT assay (n=3) with MDA-MB-231 cells treated with indicated epigenetic drug in combination of 4μM NVP-ADW742 for 72 hours. (D) RT-qPCR analysis of *FYN* mRNA level in MDA-MB-231 cells treated with 4μM NVP-ADW742 and (left) 20μM pinometostat, or (right) 10μM GSK-J4 for 48 hours (n=3). (E) RT-qPCR analysis of MDA-MB-231 Cas9 cells expressing indicated sgRNAs and treated with 4μM NVP-ADW742 for 48 hours. (F) Western blot analysis of MDA-MB-231 Cas9 cells expressing indicated sgRNAs and treated with 4μM NVP-ADW742 for 48 hours. (G) RT-qPCR analysis of FYN mRNA level in MDA-MB-231 Cas9 cells expressing indicated sgRNAs treated with NVP-ADW742 (n=4). (H) RT-qPCR analysis of KDM4 mRNA levels in MDA-MB-231 cells treated with gefitinib (n=3). (I) RT-qPCR analysis of indicated genes in TNBC primary tumor derived organoids treated with indicated TKIs (n=1). All data are plotted as mean±s.d. Unpaired two-sided Student's t-test in C and H.*, $p<0.05$; **, $p<0.01$; ***, $p<0.001$; n.s., $p>0.05$. All replicates are biological replicates.

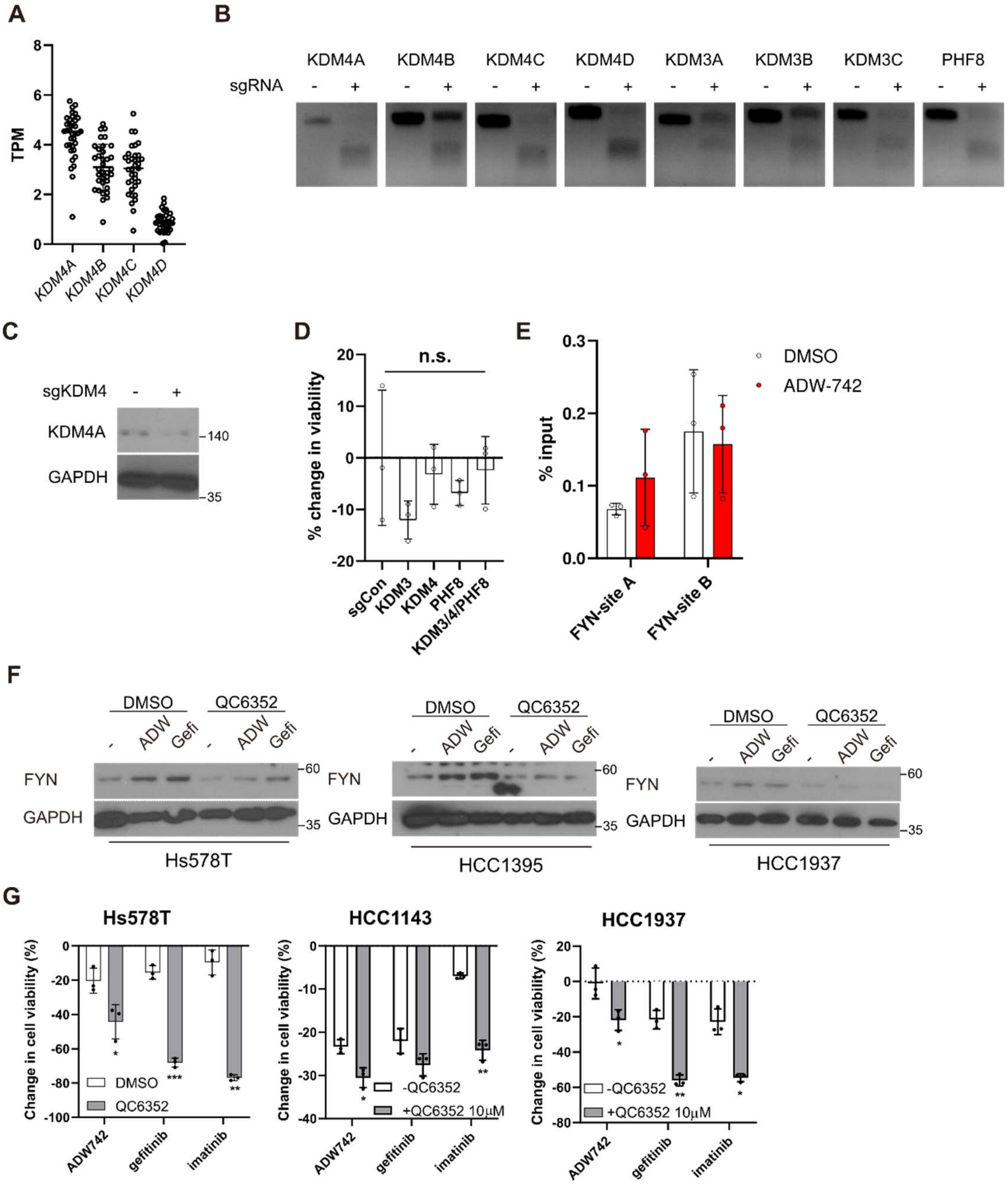


**Figure S6. KDM4 inhibition synergizes with TKIs in multiple TNBC cell lines.** (A) expression levels of KDM4 family genes in TNBC cell lines in CCLE. (B) T7 endonuclease assay validation of sgRNAs against indicated genes used in figure 3I. (C) Validation of KDM4 knockout by western blot in MDA-MB-231 cells expressing indicated sgRNA. (D) MTT assay with MDA-MB-231 Cas9 cells expressing indicated sgRNAs. sgCCR5 is used as a control sgRNA that is expected to minimally affect cell viability. (E) H3K27me3 Chromatin immunoprecipitation-qPCR analysis of MDA-MB-231 cells treated with 4μM NVP-ADW742 for 48 hours at specified genomic loci (n=3). (F) western blot analysis of FYN expression level in indicated TNBC cell lines treated with QC6352 in combination with TKIs. (G) Summary of MTT assay with cancer cell lines treated with indicated drugs for 72 hours (n=3). TKIs are used at concentration of 4μM, 10μM, 10μM for NVP-ADW742, gefitinib, and imatinib, respectively, unless otherwise indicated. All data are plotted as mean±s.d. Unpaired two-sided Student's t-test in C.*, $p<0.05$; **, $p<0.01$; ***, $p<0.001$; n.s., $p>0.05$. All replicates are biological replicates.

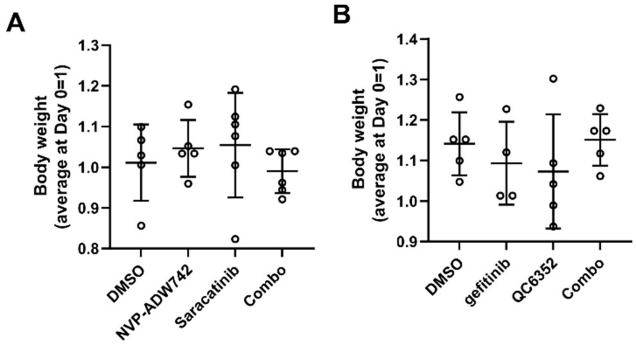


**Figure S7. No overt toxicity with drug combinations in xenograft experiments.** (A-B) Body weight of the mice described in (A) Figure 4A and (B) Figure 4B treated with indicated drug combinations (n=5-6). All data are plotted as mean±s.d. All replicates are biological replicates.

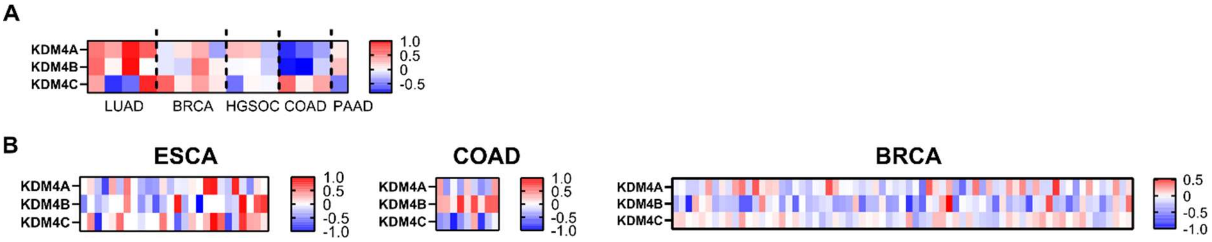


**Figure S8. KDM4 expression levels in drug tolerant cancers and residual tumor after therapy.** (A) Heatmap of log transformed ratio of KDM4 mRNA expression of drug tolerant persisters and parental cells the datasets used are identical to those used in figure 5B and 5H. (B) Heatmap of log transformed ratio of KDM4 mRNA expression of residual tumor after therapy and primary tumor before therapy. The datasets used are identical to those used in figure 5I.

**Table S1. Tyrosine kinases subject to CRISPR screens in this study.**

| | | |
|---|---|---|
| BTK | MERTK | EPHA5 |
| KDR | TYRO3 | EPHA6 |
| ABL1 | FRK | EPHA7 |
| ALK | FYN | EPHA8 |
| EGFR | HCK | EPHB1 |
| FGFR2 | SRC | EPHB2 |
| FGFR3 | TXK | EPHB3 |
| IGF1R | CSF1R | EPHB6 |
| JAK2 | INSR | TNK1 |
| JAK3 | BLK | TNK2 |
| MET | EPHA2 | ABL2 |
| FLT3 | FGR | PTK6 |
| KIT | ROS1 | PTK2 |
| PDGFRB | TYK2 | ERBB2 |
| CSK | BMX | ERBB3 |
| RET | DDR1 | ERBB4 |
| YES1 | WEE1 | INSRR |
| FLT1 | EPHB4 | LCK |
| FLT4 | ITK | PTK2B |
| JAK1 | FES | MST1R |
| FGFR1 | DDR2 | SRMS |
| FGFR4 | EPHA1 | TEK |
| LYN | EPHA10 | NTRK1 |
| SYK | EPHA3 | NTRK2 |
| PDGFRA | EPHA4 | NTRK3 |
| AXL | | |

**Table S2. List of GEO data used for analysis**

| Figure | Disease | Treatment | GEO accession number |
|---|---|---|---|
| 5B | LUAD | osimertinib | GSE193258 |
| 5H | BRCA | Lapatinib | GSE155341 |
| | COAD | Irinotecan | GSE145356 |
| | PAAD | gemcitabine | GSE189764 |
| | HGSOC | carboplatin | GSE198701 |
| 5I | BRCA | Epirubicin+docetaxel+bevacizumab | GSE87455 |
| | ESCA | CROSS+atezolizumab | GSE165252 |
| | COAD | radiotherapy | GSE15781 |

**Table S3. List of sgRNA used in the study**

| Target gene | Sequence |
|---|---|
| GFP (neg. con) | GGGCGAGGAGCTGTTCACCG |
| CCR5 (neg. con) | CATTAAAGATAGTCATCTTG |
| IGF1R | GGTACAATGTGAAAGGCCGA |
| EGFR | TGTCACCACATAATTACCTG |
| FYN #1 | TGGATACTACATTACCACCCs |
| FYN #2 | TGTGACAGGGAACTACTAGG |
| FYN #3 | GTCCCCCGAATCATTCCTTG |
| BTK | TATGAGTATGACTTTGAACG |
| ITK | AACTATCACCAACATAATGG |
| PTK6 | CGCACCCGACAGGACGTAGT |
| PDGFRA | AAATAATCCGTCATTCCTAG |
| KDM5A | GCTGGGATTCAAATAACTCG |
| KDM5B | TCTTGCAGATCATCTCATCG |
| KDM6A #1 | CAATTGTCAGAAGTATTCTG |
| KDM6A #2 | CTAGCAATTCAGTAACACAG |
| KDM6B #1 | GGTGCTAGAAGAGATCAGCC |
| KDM6B #2 | GCAGTCGGAAACCGTTCTTG |
| KDM4A | TGTGCACAGTTATGCCAAAG |
| KDM4B | TCACCAGGTACTGTACCCCG |
| KDM4C | GCAAGAGTATAATGCAACAG |
| KDM4D | AAATCGGTGAATTATAGATG |
| KDM3A | CATTCTGTAAGAGCGAAATG |
| KDM3B | GGATGGGTCATGCATCAATG |
| KDM3C | TTGGCATTACATCACGACGC |
| PHF8 | AGGGGCATGATACACACAAG |

**Dataset S1 (separate file).** Growth phenotype scores Z and gene interaction scores GI calculated with combinatorial screens in figure 1.

**Dataset S2** (separate file). Raw viability data for syngergyfinder analyses in figures 2F, 3J and S4B
